# Long-term metabolic consequences of acute dioxin exposure differ between male and female mice

**DOI:** 10.1101/762476

**Authors:** Myriam P Hoyeck, Hannah Blair, Muna Ibrahim, Shivani Solanki, Mariam Elsawy, Arina Prakash, Kayleigh RC Rick, Geronimo Matteo, Shannon O’Dwyer, Jennifer E Bruin

**Author notes:** Address correspondence to: Dr. Jennifer Bruin, 244 Nesbitt Biology Building, 1125 Colonel By Drive, Ottawa, ON K1S 5B6.

## Abstract

Exposure to environmental pollutants is consistently associated with increased diabetes risk in humans. In male mice, acute dioxin (2,3,7,8-tetrachlorodibenzo-*p*-dioxin; TCDD) exposure supresses insulin secretion. This study investigated the long-term effects of a single TCDD injection (20 µg/kg) on glucose metabolism and beta cell function in male and female mice. TCDD-exposed males displayed modest fasting hypoglycemia for ∼4 weeks post-injection, reduced fasting insulin levels for up to 6 weeks, increased insulin sensitivity, and decreased beta cell area. TCDD-exposed females also had long-term suppressed basal plasma insulin levels, and abnormal insulin secretion for up to 6 weeks. Unlike males, TCDD did not impact insulin sensitivity or beta cell area in females, but did cause transient glucose intolerance 4 weeks post-exposure. Our results show that a single exposure to dioxin can supress basal insulin levels long-term in both sexes, but effects on metabolism are sex-dependent.

## INTRODUCTION

Diabetes mellitus is one of the leading causes of death worldwide, and its prevalence is increasing at alarming rates (WHO, 2018). Type 2 diabetes (T2D) is the most common form of diabetes, and is defined by chronic hyperglycemia induced by peripheral insulin resistance and insufficient insulin production by pancreatic beta cells (Porte, 1991). The rapid increase in diabetes incidence worldwide cannot be accounted for by genetics and lifestyle alone, and mounting research suggests that environmental pollutants may contribute to diabetes etiology. In particular, epidemiological studies have consistently shown a correlation between exposure to persistent organic pollutants (POPs) and diabetes incidence (Sun et al., 2017; Taylor et al., 2013).

POPs are lipophilic chemicals that are highly resistant to degradation, leading to widespread global dispersion, bioaccumulation, and long-term release into the environment (Breivik et al., 2004). Dioxins and dioxin-like compounds are a broad class of POPs that accumulate in the liver and activate the aryl hydrocarbon receptor (AhR), which induces upregulation of cytochrome P450 *(Cyp)1a1* and *Cyp1a2* target genes. CYP enzymes are essential for the metabolism and excretion of xenobiotic compounds. However, CYP-mediated metabolism generates intermediate metabolites that are often more toxic than the parent compound and can cause oxidative stress and DNA/protein damage (Mimura and Fujii-Kuriyama, 2003). Our lab has shown that CYP1A enzymes are inducible and functional in human and mouse pancreatic islets, and that these enzymes remain active for at least two weeks following a single acute exposure to the highly persistent dioxin, 2,3,7,8-tetrachlorodibenzo-*p*-dioxin (TCDD) (Ibrahim *et al*., 2019). These data indicate that dioxins reach the endocrine pancreas *in vivo*, which may have important implications for islet cell physiology.

Mounting evidence suggests that dioxin exposure impairs insulin secretion, a key hallmark in diabetes progression. For example, an association between increased serum POP concentrations, including dioxin-like polychlorinated biphenyls (PCBs), and decreased insulin secretion was reported in humans (Lee et al., 2017). Suppressed insulin secretion was also observed in rodent beta cell lines after direct exposure to organochlorine pesticides (OCPs) and PCBs *in vitro* (Lee et al, 2017; Mailloux et al 2015), and in isolated rat and mouse islets 24-hours following *in vivo* exposure to TCDD (Novelli et al., 2005). Our lab has also shown that direct exposure of mouse and human islets to TCDD for 48-hours *ex vivo* causes reduced insulin secretion (Ibrahim *et al*., 2019). Furthermore, a single high-dose injection of TCDD (200 µg/kg) in male mice led to reduced plasma insulin levels for 2 weeks *in vivo*, and impaired glucose-stimulated insulin secretion by isolated islets *ex vivo* after 1 week. We also observed a substantial increase in beta cell apoptosis in these mice, suggesting that TCDD affects both beta cell function and survival (Ibrahim *et al*., 2019). However, this high dose of TCDD caused significant weight loss and severe hypoglycemia after 2 weeks, which prevented us from assessing longer-term effects of transient TCDD exposure *in vivo*.

Another limitation of our previous study was that we only examined male mice, similar to other groups (Gorski and Rozman, 1987; Gorski et al., 1988; Kurita et al., 2009; Mailloux et al., 2015). To our knowledge, sex differences in metabolic outcomes following dioxin exposure have only been reported in epidemiological studies, which suggest that women may be more susceptible than males to dioxin-induced diabetes (Consonni et al., 2008; Wang et al., 2008; Warner et al., 2013). Therefore, males and females may respond differently to pollutant exposure, and studies in males should not be used to predict the effects of pollutants in females. The purpose of this study was to investigate the long-term implications of a single exposure to TCDD on glucose homeostasis and islet cell physiology in male and female mice, using a lower dose than in our previous work.

## METHODS

### Animals

All mice received *ad libitum* access to a standard irradiated diet (Harlan Laboratories, Teklad Diet #2918, Madison, WI) and were maintained on a 12 hour light/dark cycle throughout the study. All experiments were approved by the University of British Columbia or Carleton University Animal Care Committees and carried out in accordance with the Canadian Council on Animal Care guidelines.

#### Cohort 1 (Figure 1)

As outlined in **Figure 1A**, 8 week old male C57Bl/6 mice (Jackson Labs) received a single i.p. injection of corn oil (25 ml/kg, vehicle control; n = 13), 20 μg/kg TCDD (n = 13), or 200 μg/kg TCDD (n = 4). Islets were isolated from a subset of mice on days 14 (n = 4 per group) and 28 (n = 4 per group; corn oil and 20 μg/kg TCDD mice only) for RNA isolation and Cyp1a1 enzyme activity assays. Islets for RNA were stored in RLT buffer (Qiagen) at −80°C. Whole pancreas and liver were harvested from a different subset of mice on day 28 (n = 5 per group; corn oil and 20 μg/kg TCDD mice only) and stored in 4% paraformaldehyde (PFA) for 24 hours, followed by long-term storage in 70% ethanol. A piece of liver was also flash frozen in liquid nitrogen and stored at −80°C for RNA isolation.

**Figure 1:**
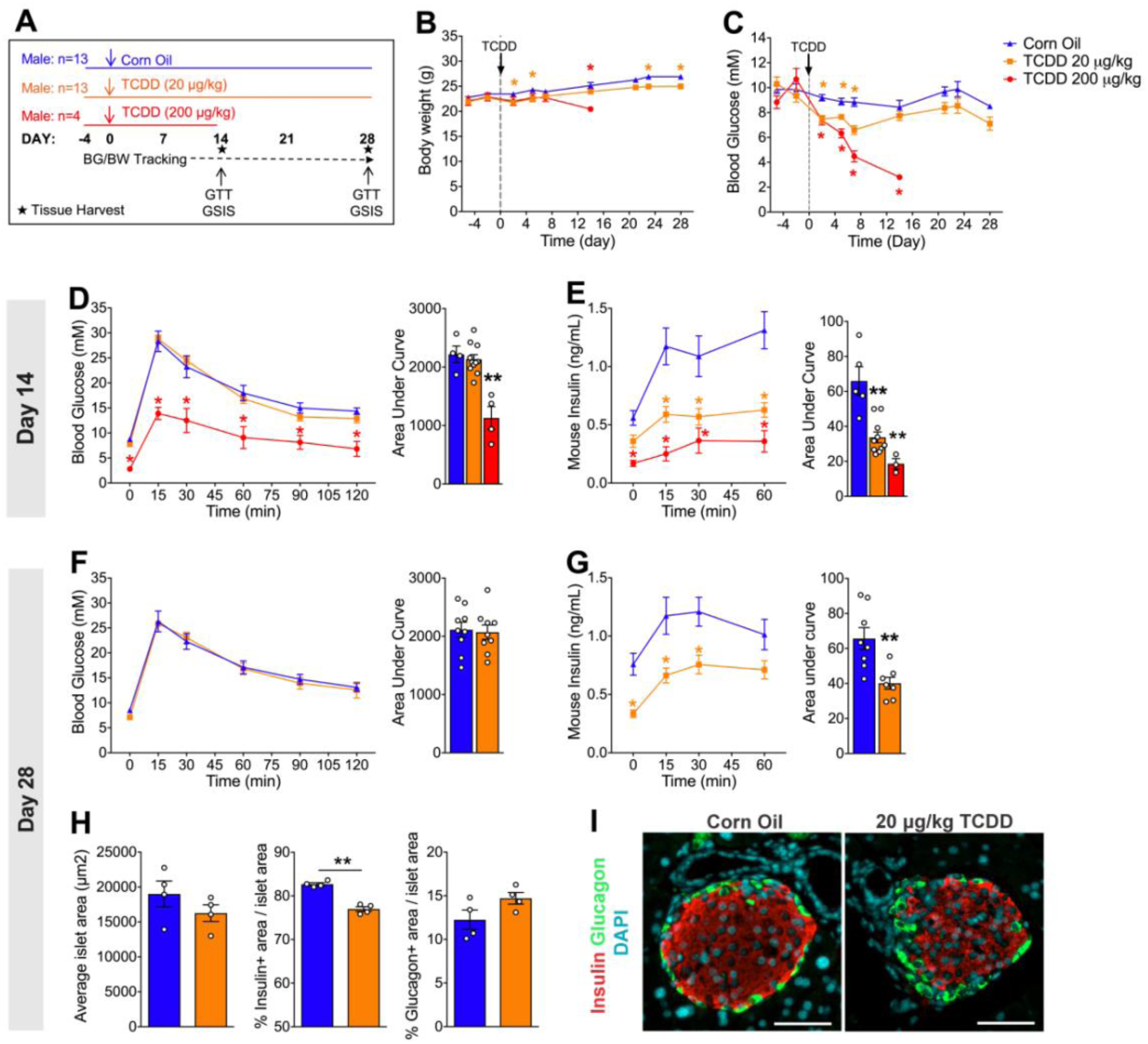
Long-term reduction in plasma insulin levels, but normal glucose tolerance *in vivo* after a single dose of TCDD in male mice. **(A)** Male mice were injected with either corn oil, 20 μg/kg TCDD or 200 μg/kg TCDD on day 0 and followed for up to 4 weeks. BG = blood glucose; BW = body weight; GTT = glucose tolerance test; GSIS = glucose-stimulated insulin secretion. **(B)** Body weight and **(C)** blood glucose levels were measured after a morning fast throughout the study. **(D-F)** Blood glucose and **(E-G)** plasma insulin levels were measured during an i.p. GTT on day 14 **(D-E)** and day 28 **(F-G). (H)** Average islet area, and % of islet area that is immunoreactive for either insulin or glucagon. **(I)** Representative images of paraffin-embedded pancreas sections on day 28 after exposure to control or TCDD (20 μg/kg). Tissues are stained with insulin (red), glucagon (green), and DAPI (blue). Scale bars = 50 μm. All data are presented as mean ± SEM. Individual data points on bar graphs represent biological replicates (different mice). *p<0.05, **p<0.01 versus control. **(B-C)** Day 5-14: two-way ANOVA with Dunnett test for multiple comparisons. Day 21-28: two-way ANOVA with Sidak test for multiple comparisons. **(D-E)** Line graphs: two-way RM ANOVA with Dunnett test. AUC: one-way ANOVA with Dunnett test. **(F-G)** Line graphs: two-way RM ANOVA with Sidak test. AUC: two-tailed unpaired t-test. **(H)** Unpaired two-tailed t-test.

#### Cohort 2 (Figures 2-4)

As outlined in **Figure 2A**, 8 week old male (n = 12) and female (n = 14) C57Bl/6 mice (bred in-house at the Modified Barrier Facility, University of British Columbia) received a single i.p. injection of corn oil (25 ml/kg, vehicle control) or 20 μg/kg TCDD. All mice were euthanized on day 42 to harvest pancreas in 4% PFA for histological analysis.

**Figure 2:**
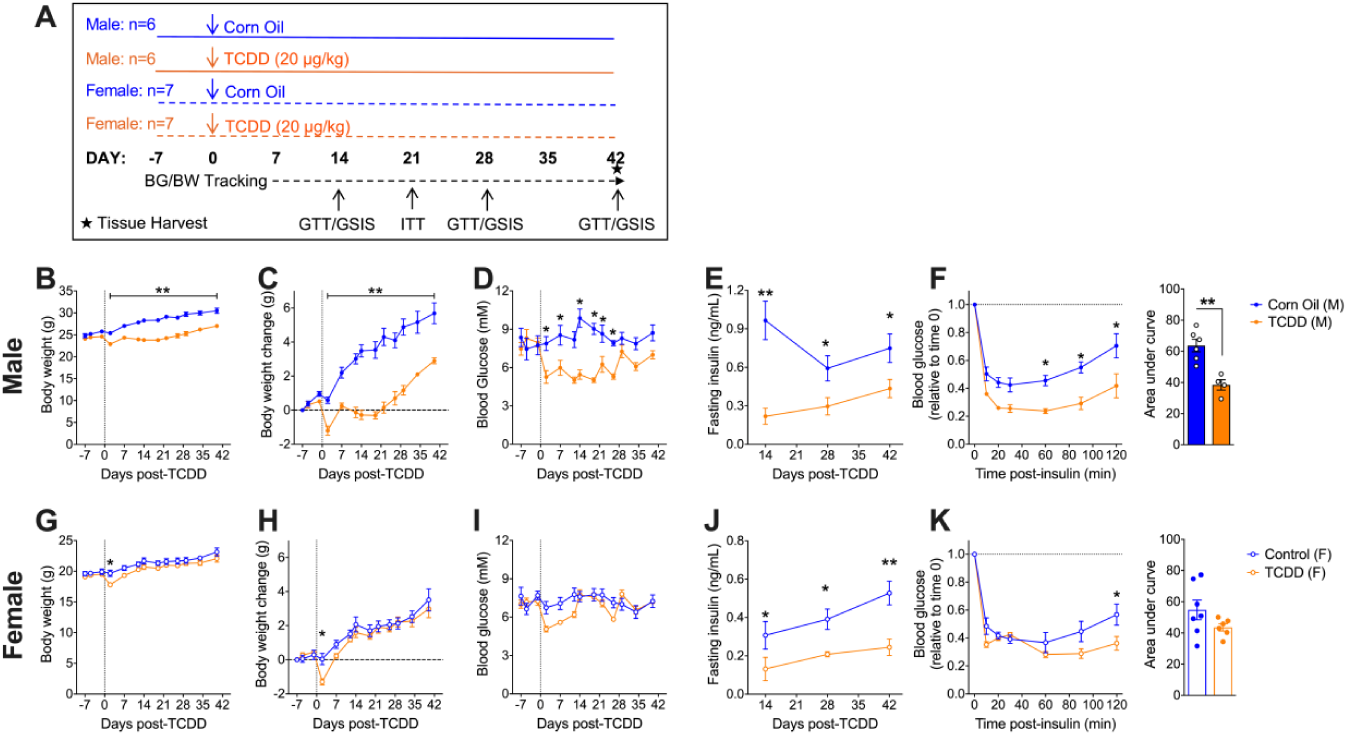
Sex differences in fasting metabolic response and insulin sensitivity following TCDD exposure. **(A)** Male and female mice were injected with either corn oil or 20 μg/kg TCDD on day 0 and followed for up to 6 weeks. BG = blood glucose; BW = body weight; ITT = insulin tolerance test; GTT = glucose tolerance test; GSIS = glucose-stimulated insulin secretion. **(B-C,G-H)** Body weight was measured throughout the study in males **(B-C)**, and females **(G-H)**. Data are presented both as raw values **(B,G)**, and change in body weight relative to the first measurement on day 7 **(C,H). (D,I)** Blood glucose and **(E,J)** plasma insulin levels were measured after a 4 hour morning fast throughout the study in males **(D-E)**, and females **(I-J). (F,K)** Blood glucose levels during an ITT on day 21 (values are normalized relative time 0 for each mouse). All data are presented as mean ± SEM. Individual data points on bar graphs represent biological replicates (different mice). *p<0.05, **p<0.01 versus control. **(B-D,G-I)** Two-way ANOVA with Sidak test for multiple comparisons. **(E,J)** Two-way ANOVA with Fisher LSD test for multiple comparisons. **(F,K)** Line graphs: two-way RM-ANOVA with Sidak test for multiple comparisons; AUC: unpaired two-tailed t-test.

**Figure 3:**
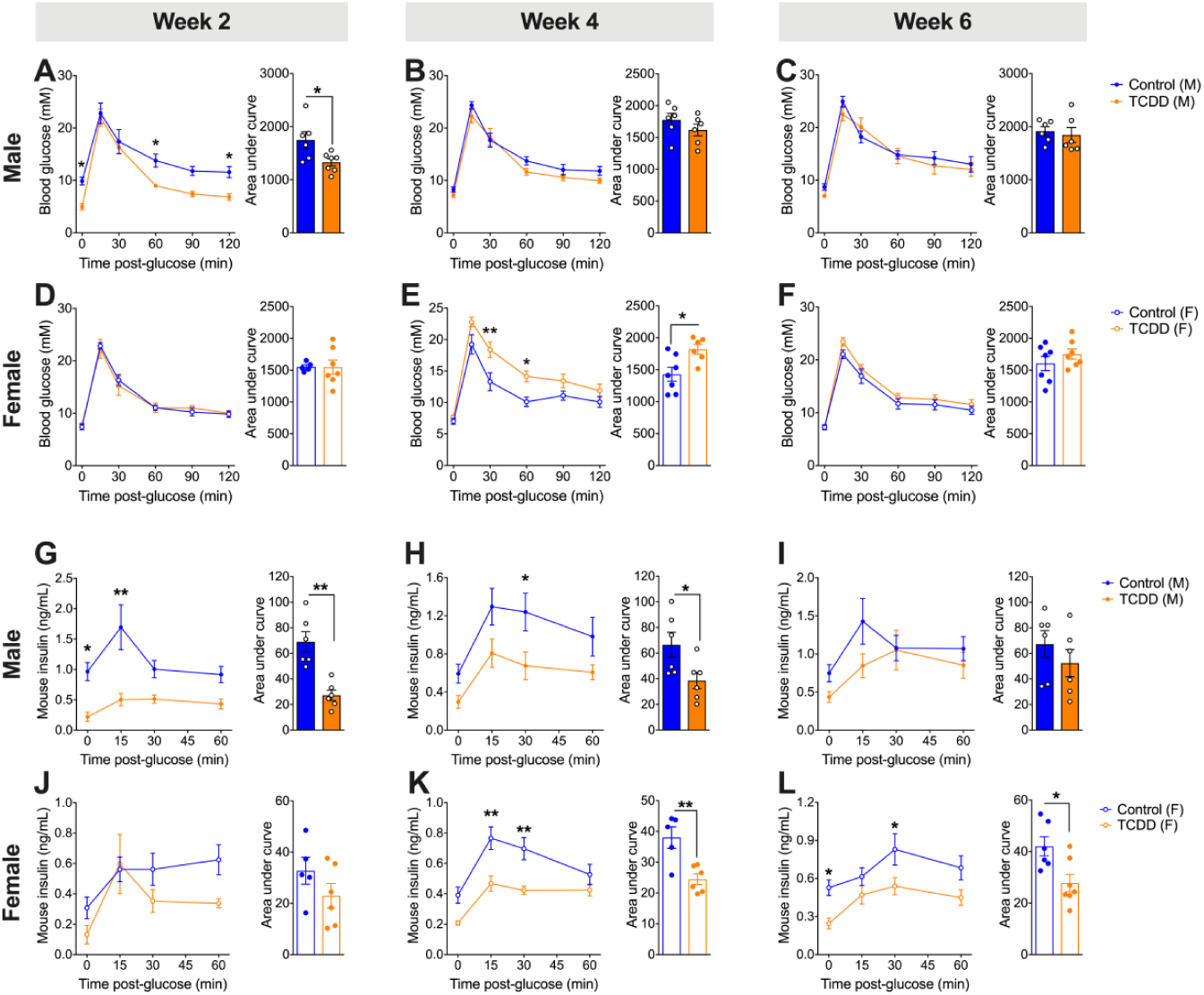
Transient TCDD exposure leads to long-term suppression of plasma insulin in both sexes, but sex differences in overall glucose tolerance *in vivo*. **(A-C, G-I)** Male and **(D-F,J-K)** female mice were injected with either corn oil or 20 μg/kg TCDD on day 0 and glucose tolerance and glucose-stimulated insulin secretion was assessed *in vivo* on days 14, 28, and 42 (see Figure 2A for schematic timeline). **(A-F)** Blood glucose levels during a GTT on days 14 **(A,D)**, 28 **(B,E)**, and 42 **(C,F). (G-L)** Plasma insulin levels during the GTT test on days 14 **(G,J)**, 28 **(H,K)**, and 42 **(I,L).** All data are presented as mean ± SEM. Individual data points on bar graphs represent biological replicates (different mice). *p<0.05, **p<0.01 versus control. **(A-L)** Line graphs: two-way RM-ANOVA with Sidak test for multiple comparisons; AUC: unpaired two-tailed t-test. See also Figure S2 for additional data from a second cohort.

**Figure 4:**
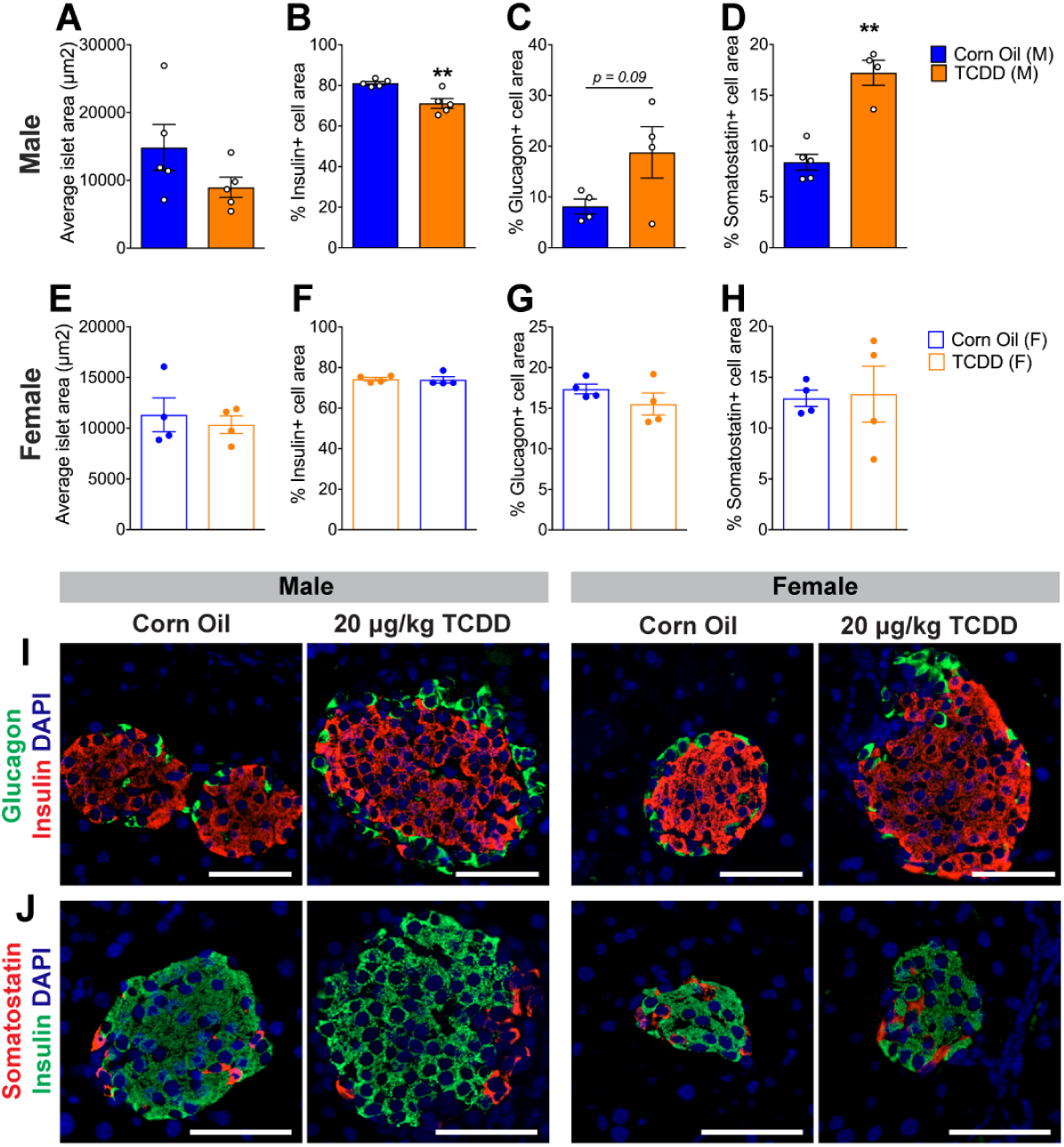
TCDD exposure decreases % beta cell area and increases % delta cell area in male mice but does not affect islet cell composition in female mice. **(A-D)** Male and **(E-H)** female mice were injected with either corn oil or 20 μg/kg TCDD on day 0 and pancreas tissue was harvested on day 42 for analysis by immunofluorescence staining (see Figure 2A for schematic timeline). **(A-B)** Average islet area. **(B-D,F-H)** % of islet area that is immunoreactive for insulin (**B,F)**, glucagon **(C,G)**, or somatostatin **(D,H). (I)** Representative images of pancreas sections from male and female mice exposed to either corn oil or TCDD showing immunofluorescence staining for insulin/glucagon or insulin/somatostatin. Scale bars = 50 μm. All data are presented as mean ± SEM. Individual data points on bar graphs represent biological replicates (different mice). **p<0.01 versus control; unpaired two-tailed t-test.

#### Cohort 3 (Figure S2)

Male (n = 10) and female (n = 10) 8 week old C57Bl/6 mice (Charles River) received a single i.p. injection of corn oil (25 ml/kg, vehicle control) or 20 μg/kg TCDD. All mice were euthanized after 28 days.

### In Vivo Metabolic Assessments

All metabolic analyses were performed in conscious, restrained mice and blood samples were collected via saphenous vein using heparinized microhematocrit tubes at the indicated time points. Blood glucose levels were measured using a handheld glucometer (Lifescan; Burnaby, Canada).

Body weight and blood glucose levels were assessed weekly or bi-weekly following a 4-hour morning fast. For all other metabolic tests, time 0 indicates the blood sample collected after fasting and prior to administration of glucose or insulin. For glucose tolerance tests (GTTs), mice received an i.p. bolus of glucose (2 g/kg; Vétoquinol) following a 6-hour morning fast. During the GTT, blood was collected at the indicated time points for measuring plasma insulin levels by ELISA (ALPCO mouse ultrasensitive insulin ELISA, #80-INSMSU-E01). For insulin tolerance tests (ITTs), mice received an i.p. bolus of insulin (0.7 IU/kg, Novolin ge Toronto; Novo Nordisk Canada #02024233) after a 4-hour morning fast.

### Mouse Islet Isolation & Culture

Islets were isolated from mice by pancreatic duct injection with collagenase (1,000 units/ml; Sigma Aldrich #C7657) dissolved in Hanks’ balanced salt solution (HBSS: 137 mM NaCl, 5.4 mM KCl, 4.2 mM NaH_2_PO_4_, 4.1 mM KH_2_PO_4_, 10 mM HEPES, 1 mM MgCl_2_, 5 mM dextrose, pH 7.2). Pancreata were incubated at 37°C for 12 minutes, vigorously agitated, and the collagenase reaction quenched by adding cold HBSS with 1mM CaCl_2_. The pancreas tissue was washed 3-times in HBSS+CaCl_2_ (centrifuging for 1 min at 1000 rpm in between washes) and resuspended in Ham’s F-10 (HyClone #SH30025.01 or Corning #10-070-CV) containing 0.5% bovine serum albumin (BSA; Sigma-Aldrich #10775835001), 100 units/ml penicillin, and 100 μg/ml streptomycin (Corning #30002CI). Pancreas tissue was filtered through a 70 μm cell strainer and islets were handpicked under a dissecting scope to >95% purity.

### Cytochrome P450 Enzyme Activity Assay

Enzyme activity was measured in isolated mouse islets (50 islets per well) using the Promega P450-Glo™ CYP1A1 Assay (#V8752) in 96-well white-walled plates (Greiner Bio-One, # 655098) using the lytic method, as described by the manufacturer.

### Quantitative Real Time PCR

RNA was isolated from liver samples using the Qiagen RNeasy Mini Kit (#74104) or Trizol™ Reagent (Invitrogen #15596018), and mouse islets using the Qiagen RNeasy Micro Kit (#74004). DNase treatment was performed prior to cDNA synthesis with the iScript™ gDNA Clear cDNA Synthesis Kit (#1725035, Biorad, Mississauga, ON Canada). Quantitative real-time PCR (qPCR) was performed using the SsoFast™ EvaGreen Supermix (#1725200, Biorad) or SsoAdvanced™ Universal SYBR^®^ Green Supermix (#1725271, Biorad) and run on a CFX96 or CFX384 (Biorad). *Hprt* or *Ppia* were used as the reference genes. Data were analyzed using the ΔΔCT Method. Primer sequences are listed in **Table S1.**

### Immunofluorescent staining and image quantification

Paraffin sections (5 μm thickness) were prepared by the University of Ottawa Heart Institute Histology Core Facility (Ottawa, ON). Immunofluorescent staining was performed as previously described (Ibrahim et al 2019) and the following antibodies were used: rabbit anti-insulin (C27C9, 1:200, Cell Signaling #3014), mouse anti-insulin (L6B10, 1:250, Cell Signaling # 8138BF), mouse anti-glucagon (1:1000, Sigma-Aldrich #G2654), rabbit anti-somatostatin (1:500, Sigma-Aldrich # HPA019472). For islet quantification, a minimum of 8 islets were imaged with an EVOS FL Cell Imaging System (invitrogen). For each islet, the area was measured for the whole islet, insulin+ immunoreactivity, glucagon+ immunoreactivity, and somatostatin+ immunoreactivity using ImageJ software. The % hormone+ area was calculated as hormone+ area / islet area x 100 and the average value was calculated for each mouse.

### Statistical Analysis

All statistics were performed using GraphPad Prism software (GraphPad Software Inc., La Jolla, CA). Specific statistical tests are indicated in figures legends. Sample sizes are described in the Star Methods (EXPERIMENTAL MODEL AND SUBJECT DETAILS) and are also shown in schematic timelines (**Figure 1A, 2A**). For all analyses, p<0.05 was considered statistically significant. Data are either presented as mean ± SEM (line graphs) or as bar plots with individual data points overlaid. Individual data points always represent biological replicates (different mice).

## RESULTS

### A single dose of TCDD causes a long-term reduction of plasma insulin levels in male mice

To examine long-term effects of acute TCDD exposure on glucose homeostasis we first tested whether the previously observed decrease in plasma insulin levels following a high dose TCDD injection (200 µg/kg) could be reproduced with a lower dose model (20 µg/kg). Furthermore, it was critical to find a dose of that did not cause TCDD-induced lethality and wasting syndrome (i.e. severe weight loss and dramatic glycemia)(Uno et al., 2004) so the mice could be maintained long-term (see **Fig 1A** for study timeline). Consistent with our previous study (Ibrahim et al 2019), 200 µg/kg TCDD induced a significant decline in body weight and a dramatic decrease in blood glucose levels to ∼3 mM within 12 days following exposure (**Fig 1B-C**). However, 20 µg/kg TCDD only induced a modest decrease in body weight, and fasting blood glucose remained within a normal physiological range for the duration of the study (> 6 mM; **Fig 1B-C**). During a glucose tolerance test at 2 weeks post-TCDD, mice exposed to 200 µg/kg TCDD showed severe hypoglycemia (**Fig 1D**), which was associated with a dramatic decrease in plasma insulin levels (**Fig 1E**). Interestingly, 20 µg/kg TCDD had no effect on glycemia at either 2 or 4 weeks post-exposure (**Fig 1D,F**), but plasma insulin levels were significantly decreased throughout the glucose challenge at both time points (**Fig 1E,G**). In addition, the effect of TCDD on insulin secretion was clearly dose-dependent at 2 weeks post-exposure (**Fig 1E**). We were unable to study the mice exposed to 200 µg/kg TCDD after 2 weeks due to weight loss, but the long-term suppression of plasma insulin levels in mice exposed to 20 µg/kg TCDD was associated with a significant decrease in insulin+ area within islets at 4 weeks (**Fig 1H-I**). There was no significant change in glucagon+ area or overall islet size (**Fig 1H-I**).

We further validated the 20 µg/kg TCDD dosing model by ensuring that TCDD was reaching the endocrine pancreas, using Cyp1a1 induction as a biomarker for direct *in vivo* exposure (Ibrahim et al 2019). At 2-4 weeks following the 20 µg/kg or 200 µg/kg TCDD injection, *Cyp1a1* expression was induced ∼50-fold and ∼40-fold, respectively, in the liver (**Fig S1A**) compared to ∼3-fold and ∼12-fold, respectively, in the islets (**Fig S1B**). Moreover, we also observed a dose-dependent effect of TCDD on Cyp1a1 enzyme activity in islets (**Fig S1C**), indicating that TCDD reaches the islets and induces long-term activation of Cyp1a1 with both doses. We also verified that 20 µg/kg TCDD was not overtly cytotoxic by measuring inflammatory and apoptotic pathways in the liver and islets at 2 weeks post-TCDD. Consistent with our previous findings (Ibrahim et al 2019), we observed an increase in pro-inflammatory cytokines, *Tnfa and Il-1b*, and the anti-apoptotic gene *Birc3* in the liver (**Fig S1D**), along with a decrease in *Il-1b* and *Birc3* in the islets (**Fig S1E**) at 2 weeks following 200 µg/kg TCDD. The 20 µg/kg TCDD dose caused similar and dose-dependent changes in liver inflammation pathways (**Fig S1D**), but no changes in islets (**Fig S1E**).

### TCDD suppresses fasting insulin levels long-term in both males and females, but effects on glycemia, body weight, and insulin tolerance are sex-dependent

We next investigated whether the long-term effects of TCDD (using only the 20 µg/kg dose from this point forward) on basal metabolism and fasting insulin levels are sex dependent. Consistent with the first cohort (**Fig 1B-C**), male mice showed a sustained decrease in body weight throughout the 6-week study (**Fig 2B-C**) and reduced fasting blood glucose levels from days 2 to 26 post-TCDD (**Fig 2D**). This was associated with a long-term suppression of fasting plasma insulin levels at 2, 4, and 6 weeks post-TCDD (**Fig 2E**), and a pronounced increase in insulin sensitivity at 3 weeks (**Fig 2F**). In contrast, female mice only had a transient and very modest decrease in body weight (**Fig 2G-H**) and fasting blood glucose levels (**Fig 2I**) in the first week post-TCDD. Interestingly, TCDD-exposed females displayed long-term suppression of fasting insulin levels (**Fig 2J**), similar to males (**Fig 2E**), but with no overall change in insulin sensitivity (**Fig 2K**).

### TCDD induces sex-dependent changes in glucose tolerance and insulin secretion

To further characterize sex-dependent effects of TCDD exposure on metabolism, we assessed glucose tolerance and beta cell function for 6 weeks *in vivo* after a single TCDD injection. Male mice were modestly hypoglycemic during a glucose challenge at 2 weeks post-TCDD compared to controls, but their glycemia was restored by 4 weeks (**Fig 3A-C**). Male mice also showed a significant decrease in plasma insulin levels during GTTs at 2 and 4 weeks post-TCDD, with levels returning to normal by 6 weeks (**Fig 3G-I**). Similar trends were seen in a separate cohort of male mice, where plasma insulin levels were suppressed at 1 and 2 weeks post-TCDD (**Fig S2A-B**), with insulin levels and glucose tolerance returning to normal by 4 weeks (**Fig S2C-D**).

In contrast, female mice had normal glucose tolerance at 2 and 6 weeks post-TCDD (**Fig 3D,F)**, but were significantly hyperglycemic during a GTT at 4 weeks post-TCDD (**Fig 3E**). Importantly, this transient hyperglycemia at 4 weeks post-TCDD was replicated in a second cohort of female mice (**Fig S2H)**. Females showed normal plasma insulin levels during a glucose challenge at 2 weeks post-TCDD, but had significantly decreased insulin levels at 4 and 6 weeks (**Fig 3J-L**). In the second cohort, females were transiently hyperinsulinemic at 1 week post-TCDD, followed by reduced insulin levels at 2 weeks, and normal levels by 4 weeks (**Fig S2E-G**).

### Islet composition is altered in male but not female mice following TCDD exposure

Finally, we assessed islet composition to determine whether the decreased beta cell area observed in TCDD-exposed male mice (**Fig 1H-I**) was reproducible and whether there was sexual dimorphism in beta cell survival. Consistent with the first cohort of male mice (**Fig 1H-I)**, we again observed a modest but significant decrease (∼10%) in insulin+ area in TCDD-exposed male islets (**Fig 4B,I-J)**, a trend towards increased glucagon+ area (**Fig 4C,I**), a significant increase in somatostatin+ area (**Fig 4D,J**), and no overall change in average islet size (**Fig 4A)**. Interestingly, no changes in endocrine cell composition were observed in the females at 6 weeks after TCDD exposure (**Fig 4E-H,I-J**).

## DISCUSSION

Our study shows that a single exposure to a persistent pollutant suppresses basal insulin levels in both male and female mice for up to 6 weeks. However, overall effects on metabolism differed drastically between sexes, whereby TCDD-exposed males showed reduced beta cell mass, increased insulin sensitivity, weight loss, and hypoglycemia, while females became transiently hyperglycemic. Females also had a delayed suppression in plasma insulin levels following dioxin exposure and generally adapted differently than males to TCDD-induced metabolic changes, suggesting that females may be more susceptible to metabolism-disrupting chemicals. These data highlight the need to investigate the link between pollution and diabetes risk in both sexes, as the mechanisms of action are most likely sex-specific.

In two separate cohorts, TCDD caused a modest decrease in body weight and blood glucose in male mice throughout the 4 to 6 week study timelines. This was associated with a long-term reduction in plasma insulin levels and an increase in insulin sensitivity, but overall normal glucose tolerance. TCDD exposure also significantly altered islet composition in males by causing a decrease in beta cell area, an increase in delta cell area, and a trend towards increasing alpha cell area. Taken together, these results could point to an initial underlying insulin deficiency (caused by impaired beta cell function, decreased beta cell number, or both), which males respond to by increasing their insulin sensitivity. This leads to a period of hypoglycemia, which likely triggers the observed increase in alpha and delta cell area to restore proper glycemic equilibrium by 6 weeks post-TCDD. In our studies, TCDD-exposed male mice showed no signs of developing hyperglycemia (in fact, the opposite occurred), but the reduction in beta cell mass could be a concern for longer term diabetes susceptibility, as endogenous beta cell regeneration is extremely limited, especially in humans.

Interestingly, TCDD caused significant hyperglycemia in female mice at 4 weeks after a single exposure, indicating sexual dimorphism in diabetes susceptibility. Like males, TCDD-exposed females displayed long-term suppression of fasting insulin levels but they differed from males in all other aspects of the metabolic phenotype. TCDD caused only a transient and very modest decrease in body weight and fasting blood glucose, and no overall change in insulin sensitivity or islet composition in female mice. We also noticed that the onset of suppressed insulin secretion was consistently delayed in females compared to males, although the exact timing differed slightly between cohorts. In the first cohort TCDD decreased insulin levels during the GTT at 2-4 weeks in males versus 4-6 weeks in females; in a second cohort TCDD-exposed males were hypoinsulinemic at 1-2 weeks, whereas females were initially hyperinsulinemic at 1 week, followed by hypoinsulinemia at 2 weeks. In both cohorts females were significantly hyperglycemic during a glucose challenge at 4 weeks post-TCDD. Together these data suggest that TCDD causes insulin insufficiency in both sexes, although possibly through different mechanisms, as females did not show loss of beta cell mass. Furthermore, unlike males, female mice do not compensate for low insulin by adjusting insulin sensitivity to maintain healthy glucose homeostasis, which likely causes the abnormalities in their glycemia. The transient hyperglycemia coinciding with hypoinsulinemia are key hallmarks in diabetes progression, suggesting that females may be susceptible to diabetes following dioxin exposure.

The sex-dependent effects of TCDD on beta cell mass were quite interesting. We had previously observed that 200 µg/kg TCDD decreased expression of pro-inflammatory and anti-apoptotic genes in islets and caused a dramatic 22-fold increase in TUNEL+ beta cells at 1 week post-TCDD, suggesting a high rate of DNA damage and apoptosis in male mice (Ibrahim et al 2019). However, at 1 week post-TCDD the increase in TUNEL+ beta cells had not changed overall islet cell composition. We predicted that with more time, this increase in beta cell death would eventually lead to loss of beta cell mass. Indeed, we have now shown in two separate cohorts that a lower dose of TCDD leads to a significant reduction in % beta cell area at 4-6 weeks post-injection in male mice. Interestingly, no change in islet composition was observed in female mice, suggesting that females may be protected from TCDD-induced beta cell death. Estrogen is known to protect beta cells from injury and females have previously been reported to be protected from apoptosis-induced insulin deficiency in other models. For example, Le May *et al* (2006) demonstrated that male mice treated with the beta cell toxin streptozotocin (STZ) displayed an increase in TUNEL+ beta cells and a decrease in beta cell number per pancreas section, whereas no changes in islet architecture were observed in STZ-exposed females (Le May et al., 2006). Interestingly, when aromatase-deficient (ArKO) mice were exposed to STZ, males had similar levels of apoptosis and beta cell loss to wildtype males, whereas ArKO females displayed a dramatic decrease in beta cell number. Since aromatase is required for estradiol biosynthesis, these data suggest that estrogens protect females from toxin-induced beta cell loss. Lastly, estradiol treatment rescued STZ-induced beta cell loss, further supporting a role for estrogens in protecting beta cells from apoptosis. The potential involvement of estrogen in protecting from TCDD-induced beta cell injury remains to be investigated.

To our knowledge, this is the first report comparing sex-dependent effects of POP exposure on glucose metabolism and beta cell function in rodents. Epidemiological studies have consistently shown a link between POP exposure and diabetes incidence, however most studies have not stratified these effects according to sex (Airaksinen et al., 2011; Arrebola et al., 2013; Everett et al., 2007; Fræch et al., 2012; Gasull et al., 2012; Henriksen et al., 1997; Lee et al., 2014; Pal et al., 2013; Rignell-Hydbom et al., 2007; Wu et al., 2013). For example, an increased risk of diabetes was found in US veterans exposed to TCDD (Agent Orange) during the Vietnam war, but this study relied only on men (Henriksen et al., 1997). Other studies assessed the relationship between POP exposure and diabetes risk using both male and female participants, but did not compare the risk between sexes (Sun et al., 2017). Interestingly, a few studies have delineated the effects by sex and shown that females have a higher incidence of diabetes following POP exposure compared to males. For example, a 25 year follow up study of the Sevesco, Italy population exposed to TCDD showed that an increased risk of diabetes following exposure was limited to females (Consonni et al., 2008; Warner et al., 2013). Likewise, the consumption of rice-bran oil laced with PCBs in Yucheng, Taiwan correlated with an increase in diabetes risk in women only (Wang et al., 2008). In addition, studies on other types of pollutants (non-persistent) have reported sex-dependent effects on metabolism and diabetes incidence in humans and mice. Air pollution (including NO_2_, particulate matter, and SO_2_) was associated with an increased risk of diabetes in women but not in men (Brook et al., 2008; Sohn and Oh, 2017), whereas NO_X_ and O_3_ was associated with diabetes in both men and women, but the association was stronger in women (Renzi et al., 2018). Polycyclic aromatic hydrocarbons (PAHs) were associated with insulin resistance in women but not in men (Choi et al., 2015). Lastly, *in vivo* exposure to bisphenol (BPA), a non-persistent endocrine-disrupting contaminant, induced early onset of T1D in female NOD mice, whereas males had a delayed development of T1D (Xu et al., 2019). Taken together, these results suggest that pollutants disrupt metabolism in a sex-dependent manner and that females may be more susceptible to metabolism-disrupting chemicals than males.

Dioxin and dioxin-like pollutants have consistently been associated with an increase in diabetes risk in humans (Sun et al., 2017; Taylor et al., 2013). However, research thus far has mainly investigated the acute effects of TCDD on metabolism in male rodents. We have shown that a single transient exposure to TCDD can have long-lasting effects on beta cell function and glucose tolerance *in vivo*. Furthermore, TCDD has sex-dependent effects on metabolism, whereby males seem to have a greater ability to adapt to dioxin exposure than females. These results suggest that males and females may respond differently to pollutant exposure, emphasizing the need to study the effect of environmental pollutants on metabolism in both sexes. It will be particularly interesting to investigate whether the long-term decrease in beta cell mass in males and the transient hyperglycemia observed in females after a single high-dose of TCDD could progress into overt diabetes with an additional metabolic challenge. Further research is also required to elucidate the underlying mechanism of dioxin-induced changes in metabolism and the implications of increased diabetes risk in both sexes.

## ACKNOWLEDGEMENTS

We thank Dr. Tim Kieffer (UBC) for his mentorship and financial support of the experiments conducted in Vancouver at UBC. This research was supported by a Canadian Institutes of Health Research (CIHR) Project Grant (#PJT-2018-159590), the Canadian Foundation for Innovation John R. Evans Leaders Fund (#37231), and an Ontario Research Fund award. This work was also supported by a CIHR Foundation Grant to Dr. Timothy J Kieffer. M.P.H. was supported by an NSERC CGS Master’s award and an Ontario Graduate Scholarship (OGS). M.E. was funded by a Carleton University Dean’s Summer Research Internship. H.B. received NSERC USRA funding and a Carleton University Walker Summer Research Award.

## AUTHOR CONTRIBUTIONS

J.E.B. conceived of the experimental design. M.H. and J.E.B. wrote the manuscript. All authors were involved with acquisition, analysis, and interpretation of data. All authors contributed to manuscript revisions and approved the final version of the article.

