## Supplemental Files for "Long-term metabolic consequences of acute dioxin exposure differ between male and female mice"

**<u>Table S1</u>:** Primer sequences for qPCR

| REAGENT or RESOURCE | SOURCE | IDENTIFIER |
|---|---|---|
| Oligonucleotides | | |
| AGC TCT GAG CAC TGG AGA GA | This paper | PPIA-F |
| GCC AGG ACC TGT ATG CTT TA | This paper | PPIA-R |
| GCT GAC CTG CTG GAT TAC AT | This paper | HPRT-F |
| TTG GGG CTG TAC TGC TTA AC | This paper | HPRT-R |
| ATC ACA GAC AGC CTC ATT GAG C | This paper | Cyp1a1-F |
| AGATAGCAGTTGTGACTGTGTC | This paper | Cyp1a1-R |
| AGT CCG GGC AGG TCT ACT TT | This paper | *Tnfa*-F |
| ATG AAC ACC CAT TCC CTT CA | This paper | *Tnfa*-R |
| CTC AGG AGC AGA AGT CTG GG | This paper | *Nf-kb*-F |
| GCC GCT ATA TGC AGA GGT GT | This paper | *Nf-kb*-R |
| GCC ACC TTT TGA CAG TGA TGA G | This paper | *Il-1b*-F |
| AGC TTC TCC ACA GCC ACA AT | This paper | *Il-1b*-R |
| CCC GGA GAT CAG AGG TCA TTG | This paper | *Birc3*-F |
| GAA AGG CGC TGT CTT GAA CC | This paper | *Birc3*-R |
| TCG GGT CAG CCT CCT TAA AC | This paper | *Xiap*-F |
| TGG TGT CTG CAA GTA CAA AAG T | This paper | *Xiap*-R |

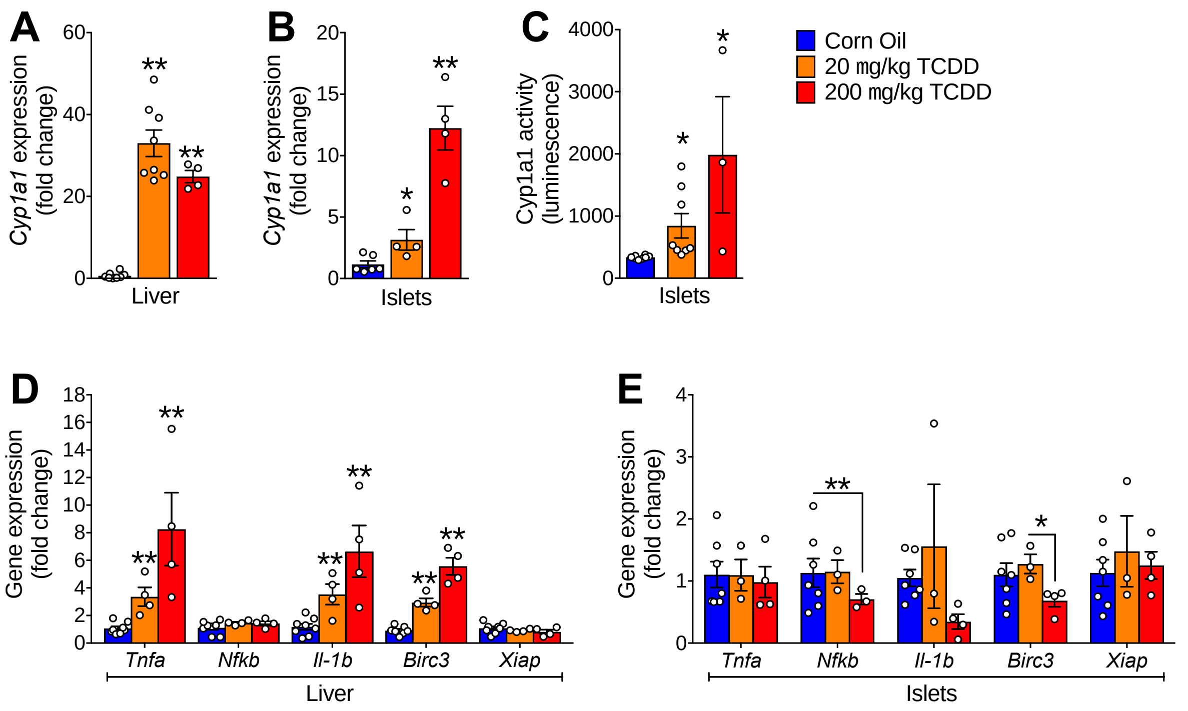


**Figure S1: Islets and liver from TCDD-exposed mice have similar induction of Cyp1a1 but different stress responses.** Male mice received a single injection of either corn oil, 20 μg/kg TCDD or 200 μg/kg TCDD. Tissues were collected either 2 or 4 weeks later. **(A-B)** *Cyp1a1* gene expression in liver **(A)** and islets **(B)**, expressed as fold change relative to control at 2-4 weeks. **(C)** Cyp1a1 enzyme activity in isolated islets at 2-4 weeks. **(D-E)** Expression of various genes related to inflammation and apoptosis at 2 weeks in liver **(D)** and isolated islets **(E)**. *p<0.05, **p<0.01 versus control unless indicated otherwise; unpaired two-tailed t-test. All data are presented as mean ± SEM and individual data points represent biological replicates (different mice).

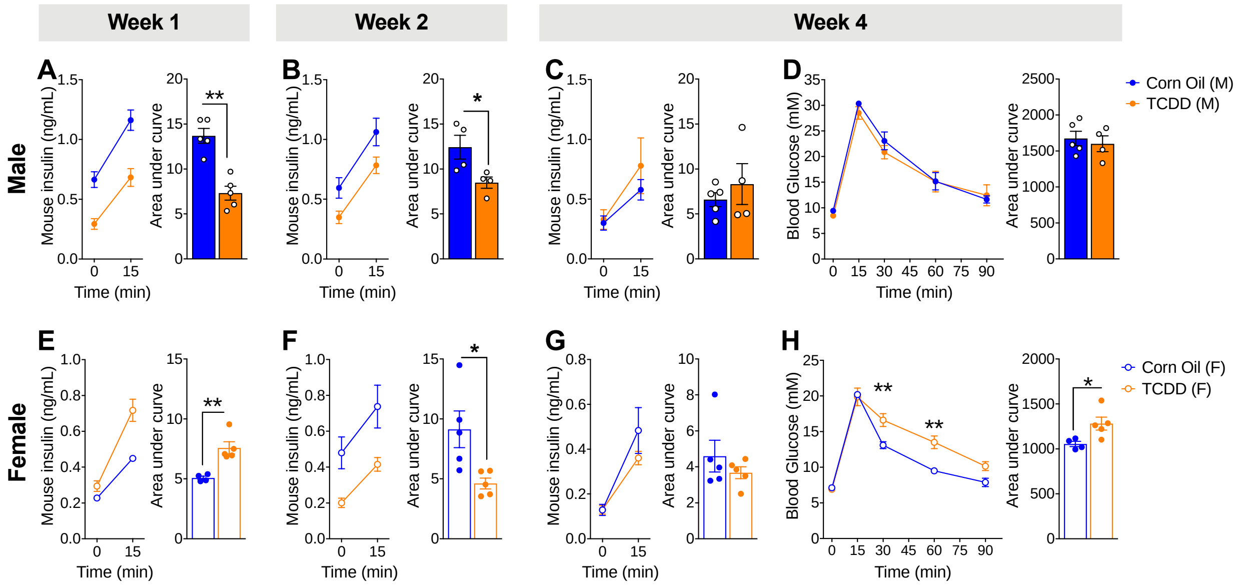


**Figure S2: Transient TCDD exposure leads to supression of plasma insulin in both sexes, but sex differences in overall glucose tolerance *in vivo*.** A new cohort of **(A-D)** male (M) and **(E-H)** female (F) mice were injected with either corn oil or 20 µg/kg TCDD on day 0 and glucose tolerance and glucose-stimulated insulin secretion was assessed *in vivo* on days 7, 14, and 28 (see Figure 3 for other cohort). **(A-C,E-G)** Plasma insulin levels before (time 0) and 15 minutes after a glucose injection on days 7 **(A,E),** 14 **(B,F),** and 28 **(C,G). (D,H)** Blood glucose levels were measured during a glucose tolerance test on day 28. All data are presented as mean ± SEM. Individual data points on bar graphs represent biological replicates (different mice). *p<0.05, **p<0.01 versus control. Line graphs: two-way RM-ANOVA with Sidak test for multiple comparisons; AUC: unpaired two-tailed t-test.
